# Salicylic acid-related ribosomal protein CaSLP improves drought and *Pst. DC3000* tolerance in pepper

**DOI:** 10.1101/2022.11.06.515320

**Authors:** Huafeng Zhang, Yingping Pei, Saeed ul Haq, Abid Khan, Rugang Chen

**Author notes:** These authors have contributed equally to this work (Huafeng Zhang, Yingping Pei). Corresponding authors: Rugang Chen, College of Horticulture, Northwest A&F University, Yangling, Shaanxi 712100, China.

## Abstract

The ribosomal protein SA plays an essential role in multiple aspects and is involved in plant growth and response to various stresses. Drought threatens pepper yield and quality. However, the resistance mechanism of pepper in response to drought are complex and not yet fully understood. Here, we describe the role of *CaSLP* in mediating pepper tolerance to drought stress. we found that *CaSLP* was highly expressed under drought and salicylic acid (SA) stress, and CaSLP was localized in cell nucleus and cytomembrane. Knockout of *CaSLP* gene significantly decreased the pepper drought tolerance, while transient expression of *CaSLP* leads to drought tolerance in pepper, and overexpression of the *CaSLP* dramatically increased the drought stress tolerance in Arabidopsis. Furthermore, exogenous spring salicylic acid enhanced drought tolerance. The characterization of resistance molecular mechanisms in the Pseudomonas syringae pv. *Tomato* DC3000 (*Pst.DC3000*) is of great significance for the pepper yield and quality, we found that *CaSLP*-knockdown pepper plants demonstrated decreased *Pst.DC3000* tolerance, whereas ectopic expression of the *CaSLP* increased the *Pst.DC3000* stress tolerance in Arabidopsis. Yeast two-hybrid (Y2H) and bimolecular fluorescence complementation (BiFC) results showed that CaNAC035 physically interacts with CaSLP in the cell nucleus, and the *CaNAC035* was identified as an upstream partner of the *CaPR1* promoter and activated the transcription. Taken together, our data demonstrated that *CaSLP* plays an essential role in the regulation of drought stress. Our study elucidates the roles of *CaSLP* response to drought stress tolerance. Furthermore, a possible regulatory model and molecular mechanisms under drought stress is proposed.

## 1. Introduction

Drought is the key factor that severely threatens plant growth and yield of field crops (Basu et al., 2016). Drought stress disturbs the physiological morphology of normal plants, which leads to stomatal closure and has detrimental effects on plant growth (Vurukonda et al., 2016). When drought exceeds for a long time, plant cells accumulate excessive reactive oxygen species (ROS), resulting in the damage of oxidative, eventually cause cell death (Baxter et al., 2013). When plants are subjected to drought, the stomatal will close which is a key role of plants response to drought, that decrease water loss of the plants leaf and it can also stops the flow of carbon dioxide, which stops photosynthesis (Katul et al., 2010). Stomatal plays an important role in modulating water loss and gas exchange (Dow and Bergmann, 2014). The closing process is influenced by the swelling degree of stomatal guard cells: the influx of ions and sucrose promotes the absorption of water, which causes the swelling of guard cells and forces the stomatal to open. Their efflux reduces the osmotic potential of the cell, which causes the cell to collapse and the stomatal to close (Kim et al., 2010; Lu et al., 1995; Pandey et al., 2007). Transcription factor *ZmNAC49* enhances drought tolerance by reducing stomatal density in maize (Xiang et al., 2021), *AtWRKY1* modulates stomatal movement in drought-stressed (Qiao et al., 2016), MADS-box factor *AGL16* negatively modulates drought stress via stomatal density in Arabidopsis (Zhao et al., 2020).

Salicylic acid (SA) is a multifunctional phytohormone involved in a variety of physiological processes and plays a key role in drought responses (Hayat et al., 2010). Exogenous-applied salicylic acid increased tolerance to various abiotic stresses mainly due to the increasing of anti-oxidative capacity (Horváth et al., 2007). Exogenous SA can also enhances the resistance under water deficit conditions and has been believed as a plant hormone to eliminate drought stress (Odjegba et al., 2012). Plant response to exogenous SA depends on variety, developmental stage, application concentration, application mode and endogenous SA level (Miura et al., 2014). Foliar applications of various plant hormones play an important role in drought-tolerance under different plant growth stages (Sohag et al., 2020). For instance, 2mM exogenous salicylic acid improve *Impatiens walleriana* drought tolerance (Dragana et al., 2020), foliar application of SA can enhance the drought tolerance in wheat by reducing the ROS accumulation (Ramadan et al., 2021), salicylic acid has been reported to enhance the drought tolerance in wheat by improving proline contents and the activity of enzymes (Sharma et al., 2017).

The ribosomal protein contained complex structures, which belong to polypeptide glycoprotein. The eukaryotic ribosome contained two unequal sub-units (60S and 40S), which are involved in plant growth and response to various stresses (Belknap et al., 1994). 40S ribosomal protein SA has a molecular weight of 67 kDa. It is expressed on different cell surface. Previous reports have found that ribosomal protein could enhance stress tolerance. For example, ribosomal protein large sub-unit genes, *RPL23A* showed an obvious increase in fresh weight and proline contents under drought and salt stresses (Belknap et al., 2017), Salicylic acid-related cotton (*Gossypium arboreum*) ribosomal protein GaRPL18 contributes to resistance to *Verticillium dahliae* (Gong et al., 2017), ribosomal protein AgRPS3aE plays a vital role in improving salt tolerance in crops (Liang et al., 2015).

In our earlier work, we characterized *CaNAC035* in pepper, which plays a crucial role in stress tolerance. In this study, we found CaNAC035 interacted with CaSLP in nucleus. We found that pepper *CaSLP* positively regulates drought tolerance. Plant stress resistance can be effectively improved by over-expressing or silencing one or more specific genes using transgenic or gene editing technology. This study provides a basis for future study of the genes and regulatory mechanisms underlying in pepper plants.

## 2. Materials and methods

### 2.1 Plant material and growth conditions

Pepper (*Capsicum annuum* L., ‘P70’) and Arabidopsis (Columbia) seeds were used and provided by Laboratory of Vegetable Plant Biotechnology and Germplasm Innovation, Northwest A&F University, Yangling, China. The pepper plants were grown at a light period of 16 h light/8 h dark22 °C light /18 °C dark cycle under the conditions of 75% relative humidity, with a light period of 16 h light/8 h dark. The Arabidopsis seeds were sterilized, and then grown on Murashige Skoog (MS) solid medium, then vernalized at 4°C for 1 day. After 7 days, the plants were placed at normal conditions (at 22 °C under 16 h light/8 h dark photoperiod).

### 2.2 RNA extraction and qRT-PCR analysis

Total RNA was extracted using RNA extraction kit (Tiangen, Beijing, China) following the manufacturer’s protocol. Synthesis of cDNA was performed using cDNA synthesis Kit (Takara). The qPCR was conducted on the Applied Biosystems instrument using the SYBR Green Master Mix, following the manufacturer’s protocol (Ding Ning). The quantitative real-time polymerase chain reaction (qRT-PCR) primers were designed by using NCBI (https://www.ncbi.nlm.nih.gov/) and the primer were shown in Table S1. The relative expression was calculated using the 2^-ΔΔCT^ method (Zhuo et al., 2013). Three biological replicates were used for this study.

### 2.3 Transient expression of *CaSLP* in pepper leaves

To generate transient expression of *CaSLP* in pepper leaves, the coding region of *CaSLP* was cloned and introduced into the pVBG2307::GFP vector, then the 35S:*CaSLP*:GFP and 35S::GFP were transformed into *Agrobacterium* strain GV3101, the method of transient expression in pepper leaves were performed as described by Huang et al., 2020. About the eight-leaf stage, using the syringe to infiltration the *Agrobacterium* into pepper plant leaves, the infiltrated leaves were harvested at 24 h and 48 h for this study.

### 2.4 Virus-induced gene silencing of *CaSLP*

To generate *CaSLP*-silenced plants, the virus-induced gene silencing method was performed. The pTRV1 and pTRV2 vector were used for this study, a 300-bp fragment of *CaSLP* was cloned and insert into the pTRV2 vector. The pTRV1, pTRV2 (control) and pTRV2:CaSLP were transformed into *Agrobacterium* strain GV3101 separately, the infection method was followed as described by Dai et al., 2018. The infection pepper plants were grown at *22°C* light/18 °C dark cycle. Twenty-eight days later, we extracted DNA and RNA for analyzing the transcript levels of *CaSLP* and also identified the positive plants using the specific primer.

### 2.5 Generation of *CaSLP*-OX in transgenic Arabidopsis plants

For Arabidopsis transformation, the full-length of *CaSLP* was cloned and combined into the pVBG2307::GFP vector, then the fusion vector of 35S:*CaSLP*:GFP was transformed into *Agrobacterium* strain GV3101, for transformation the floral dip method was used as previous reports (Clough and Bent, 1998). Then the transgene plants were screened on 1/2 MS solid medium, which contained 50 mg/ml kanamycin. We extracted DNA and RNA T3 transgenic Arabidopsis lines for analyzing the transcript levels of *CaSLP*, and also confirmed the positive plants. The T3 homozygous lines were used for this experiment and future studies.

### 2.6 Yeast two-hybrid and Bimolecular fluorescence complementation assay

The full-length of CaNAC035 was inserted into the pGBKT7 vector as bait plasmid., the full-length of CaSLP was inserted into the pGADT7 vector as prey plasmid, respectively. The recombinant vector was transformed into yeast Y2H, the strains were cultivated on SD (-Trp/-Leu), SD (-Trp/-Leu/-His/-Ade) and SD (-Trp/-Leu/-His/-Ade+X-α-gal) media for 3 days. The bimolecular fluorescence complementation (BiFC) was conducted as previously report (Choi et al., 2012).

### 2.7 Drought and *Pst. DC3000* stress tolerance assay

To analyze the drought tolerance in pepper plants: the leaves of *CaSLP*-silenced and control pepper plants were carried out with drought condition for 10 days. The leaves of pepper were collected at different stages after drought treatment. To further identify the role of *CaSLP* in transgenic Arabidopsis drought stress tolerance, 3-week-old T3 transgenic and WT lines were used in the experiments in which the T3 transgenic and WT lines were treated with 300 mM Mannitol. The samples for gene expression analyses were taken on seventh day and the phenotypic changes were observed and photographed. Samples for the stress-related genes expression analysis and physiological indexes were determined after the drought treatments. Well-watered plants were used as the positive control plants. For *Pst.DC3000* infection was performed as described by Zhu et al (2022).

### 2.8 Spraying salicylic acid treatment

In order to know the effects of exogenous salicylic acid on drought response of *CaSLP*, the *CaSLP*-silenced and control plants were exogenously supplied to 2mM salicylic acid, the plants were subjected to drought treatment. Exogenous treatment with 2mM salicylic acid was done for 2 days. We measured the growth performance, and performed the physiological index. We added Tween-20 (0.05%) for exogenous spraying salicylic acid on *CaSLP*-silenced and control plants respectively.

### 2.9 Histochemical staining and measurement of physiological traits

The malondialdehyde (MDA) content was performed essentially as previously described (Liu et al., 2006), the relative electrolyte leakage (REL) was determined based on an earlier study (Dahro et al., 2016), the water loss rate was examined according to the protocol of Zhang et al. (2022), the accumulation of H_2_O_2_ and O_2_^.-^ contents were conducted by histochemical staining of 3,3’-diaminobenzidine (DAB) and nitro-blue tetrazolium (NBT), respectively, using the method of Wang et al. (2011) O^2–^ content was examined based on the method of Ma et al. (2016), the H_2_O_2_ measurement was determined as described by Geng et al. (2018). The stomatal aperture assay was performed as described by Jiang et al. (2014).

### 2.10 Yeast one-hybrid (Y1H) assay

Y1H library screening was performed by using the Matchmaker Gold Yeast One-Hybrid Library and Screening kit (Clontech,CA, USA). The promoter fragments of *CaPR1* were inserted in the pAbAi vectors, the full-length CDS of *CaNAC035* was introduced into pGADT7 vector to perform a prey vector. Re-combinant plasmids were co-transformed into Y1H yeast strain, Y1H was carried out following the manufacturer’s manuals (Clontech, USA). The yeast strains were grown on the SD/Leu and SD/Leu/AbA media for 3d.

### 2.11 Dual luciferase and GUS activities assays

The promoter fragments of *CaPR1* were inserted into a pGreen62-SK vector. While the coding sequences of *CaNAC035* was inserted into a pGreen0800-luciferase (LUC) vector. The recombinant vectors were co-transformed into *Agrobacterium* Gv3101 and injected into four-week-old tobacco leaves (*Nicotiana benthamiana*). Transient expression was calculated by determining the ratio of luciferase (LUC) and Renilla (REN) using the Dual-Luciferase^®^ machine (Promega, WI, USA). Additionally, the GUS activities were performed as described by (Ma et al., 2021).

### 2.12 Statistical analysis

Statistical analyses were performed using one-way analysis of variance (ANOVA) tests with significant differences of P < 0.05 (*) and P < 0.01 (**) the statistical package SPSS (version 21.0, USA).

## 3 Results

### 3.1 Sub-cellular localization and expression profiles of CaSLP

In order to identify the sub-cellular localization of CaSLP, the 35S:CaSLP:GFP 35S::GFP was co-transformed into *Agrobacterium* GV3101, then infected into *N. benthamiana* leaves successfully. The 35S:CaSLP:GFP fusion protein showed obviously fluorescence signaling in the cell nucleus and cytomembrane of tobacco epidermal cells, indicating that CaSLP was localized in cell nucleus and cytomembrane (Fig. 1A). To analyze the expression pattern of *CaSLP*, the qRT-PCR was used to analyze the expression pattern, upon drought treatment, the expression of *CaSLP* was up-regulated within 6 h (Fig. 1B), after exogenous SA treatment, the expression of *CaSLP* was progressively reached to peak after 6 h (Fig. 1C). These results indicated that *CaSLP* is primarily induced by drought and SA.

**Figure 1.**
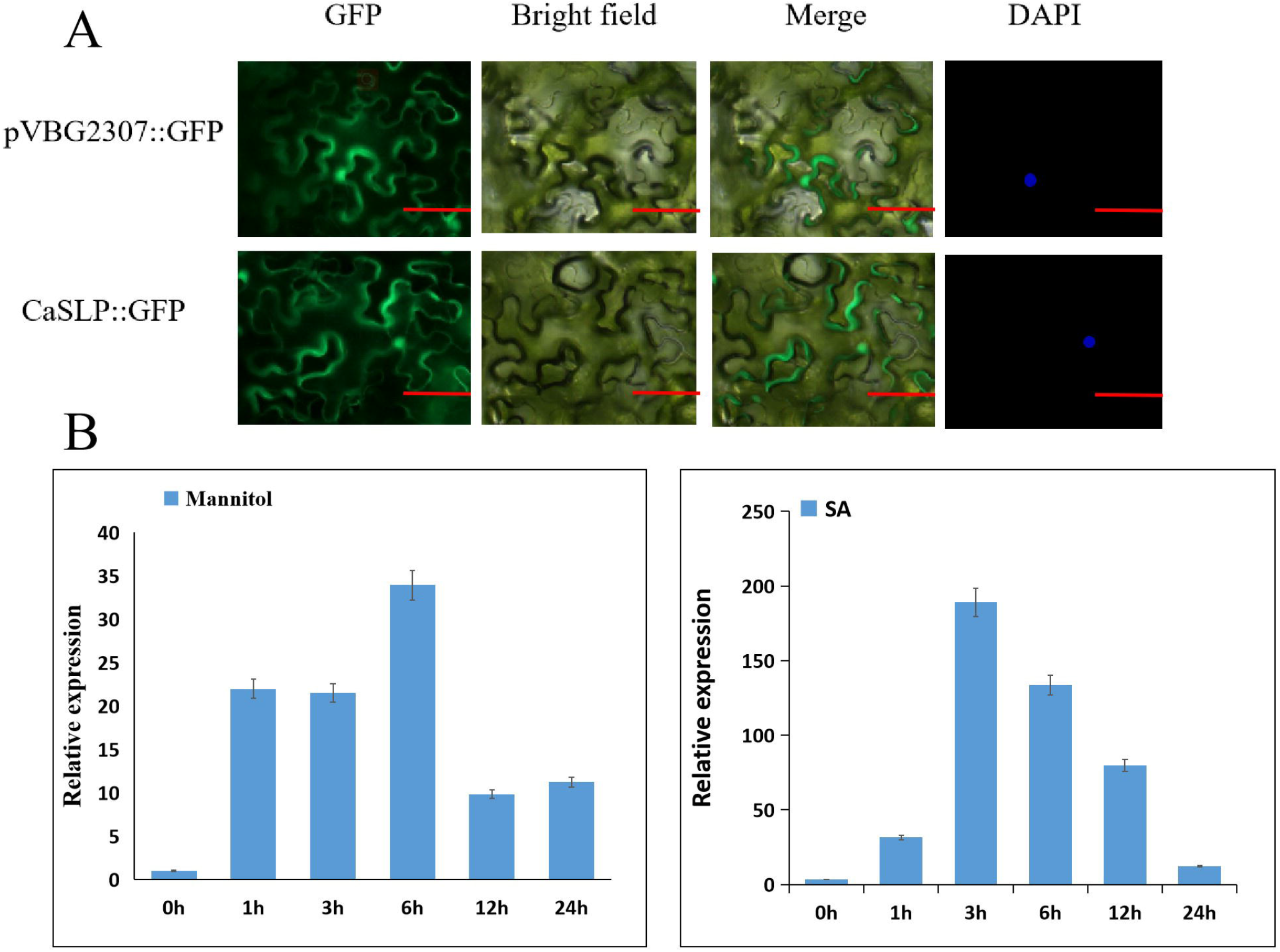
Sub-cellular localization and expression pattern assays of *CaSLP*. A, CaSLP is localized in cell nucleus and cytomembrane. B, Expression profile of *CaSLP* under drought treatment. C, Expression profile of *CaSLP* under SA hormone treatment. Scale bars = 50 μm. The data represent the means ± standard deviation (SD) (n=3, *P < 0.05, **P <0.01, Student’s t-test).

### 3.2 *CaSLP*-silenced pepper plants decreased drought tolerance

The drought-induced up-regulation of *CaSLP* implies that it has function in drought stress tolerance. To test this hypothesis, VIGS experiments were performed. As shown in the supplementary Figure S1, the plants treated with *TRV2:CaPDS* solution showed photo-bleaching in the leaves which phenotypically showed the success of the process. Then the silencing efficiency was measured through qRT-PCR, which was almost 85%. (Figure S1). No obvious morphological differences were found under normal growth conditions. Under drought stress, the *CaSLP*-silenced pepper plants showed drought-resistant phenotype compared with control pepper plants (Figure 2A). Measurement of H_2_O_2_ and O_2_^.-^ contents, the ROS indicators, the results showed that the H_2_O_2_ and O_2_^.-^ contents in the *CaSLP*-silenced pepper plants were significantly higher than control pepper plants after drought treatment (Figure 2C, F). The DAB and NBT staining of *CaSLP*-silenced pepper exhibited to stain much darker and deeper than in the control pepper plants when plants were exposed to drought stress (Figure 2B, E). When the plant were exposure to different Mannitol concentrations, the *CaSLP*-silenced pepper showed more severe leaf wilting than did the control under each Mannitol concentration (Figure 2D), and accompanying by *CaSLP*-silenced pepper plants had lower chlorophyll content than control (Figure 2G). In addition, the stomatal aperture and water loss rate also measured for this study. No apparent differences were found between the *CaSLP*-silenced and control under normal conditions. However, when subjected to drought, the VIGS plants showed slightly higher stomatal aperture and water loss rate than control plants (Figure 2H-J). All of these data demonstrated that silencing of *CaSLP* reduces drought tolerance.

**Figure 2.**
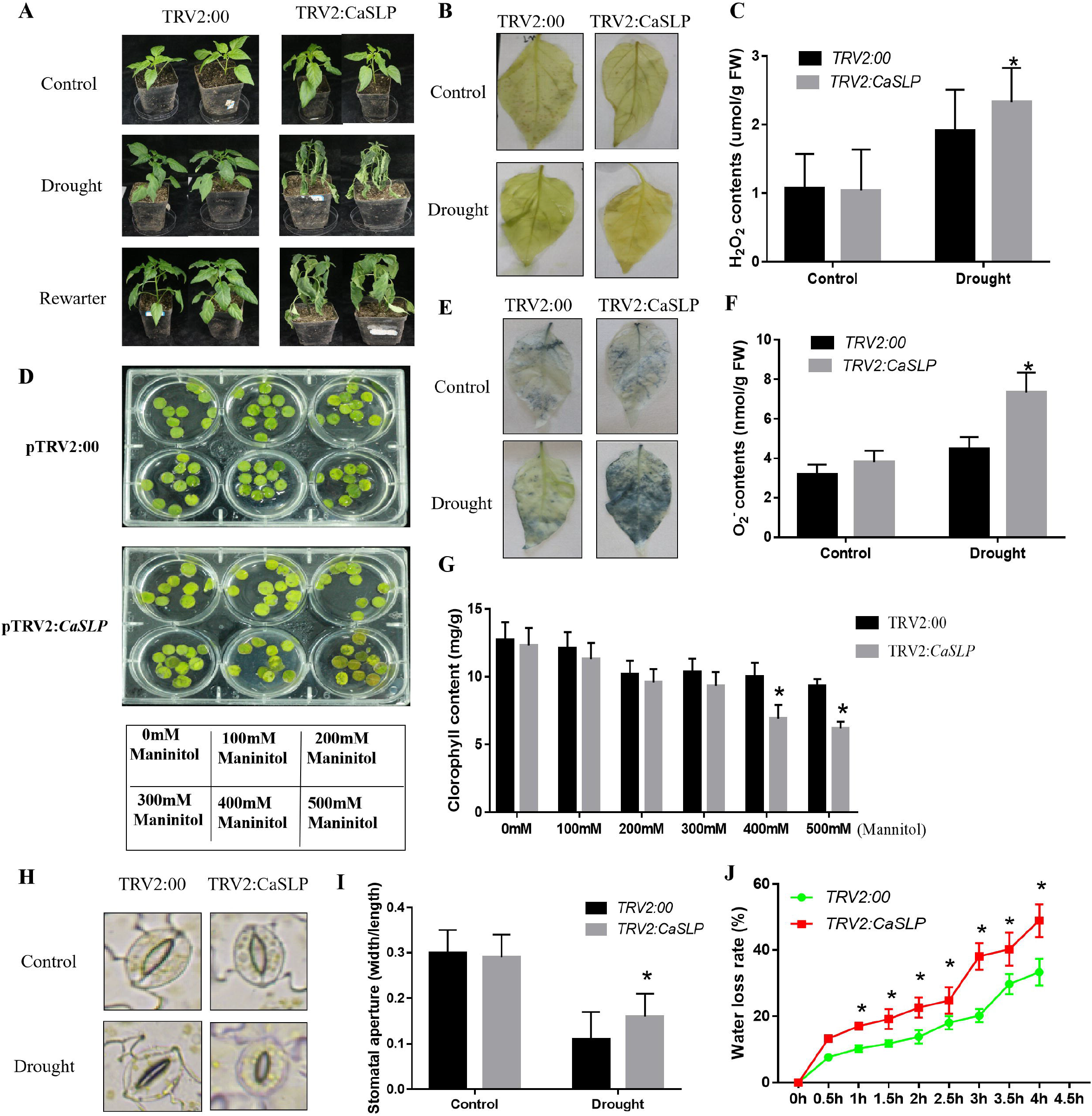
Assessment of *CaSLP*-silenced plants. A, D. The phenotypes of *CaSLP*-silenced and the control plants. B, DAB staining. C, H_2_O_2_ contents. E, NBT staining. F, O2.- contents. G, Chlorophyll contents. H, I. Stomatal aperture. J, Water loss rate. The data represent the means ± SDs (n=3, *P < 0.05, **P <0.01, Student’s t-test).

### 3.3 Transient expression of *CaSLP* improved pepper drought tolerance

To explore the function of *CaSLP* in response to drought stress tolerance, transient overexpression of *CaSLP* (*CaSLP*-To) in pepper was performed. The *CaSLP*-To and Mock plants were exposed to drought condition for 10 days. We measured the expression, and the result showed that the expression of *CaSLP*-To was visibly higher than Mock pepper plants (Fig. 3A). The *CaSLP*-To plants remained good growth and leaf turgid. Without stressful conditions, no clear differences in plants morphology between *CaSLP*-To and Mock pepper plants. However, after drought stress caused severe leaf wilting and a significant increase of ROS levels, and the H_2_O_2_ and O_2_^.-^ contents of *CaSLP*-To plants was significantly lower than that of Mock pepper plants (Fig. 3B, E, F). Next, we measured the stomatal aperture, the stomatal aperture of *CaSLP*-To and Mock plants did not differ (Fig. 3C). Whereas upon the drought exposure, the stomatal aperture of *CaSLP*-To plants showed an obviously lower stomatal aperture as compared to the control Mock pepper plants (Fig. 3G). We also measured the water loss rate, we found that the *CaSLP*-To plants displayed lower water loss rate than Mock pepper plants (Fig. 3D). These results demonstrated that the Transient expression of *CaSLP* enhanced the drought stress tolerance. Next, we measured the expression of stomatal related genes *CaSDD1, CaYOD1*, and *CaFAMA*. As shown in Figure 3H-J, the relative expression of (*CaSDD1, CaYOD1*, and *CaFAMA*) in *CaSLP*-To plants were strikingly higher than the controls pepper plants under drought stress. These results provide the key genetic evidence that induction of stomatal related genes by *CaSLP* is dependent on the *CaSLP*-mediated manner in response to drought stress.

**Figure 3.**
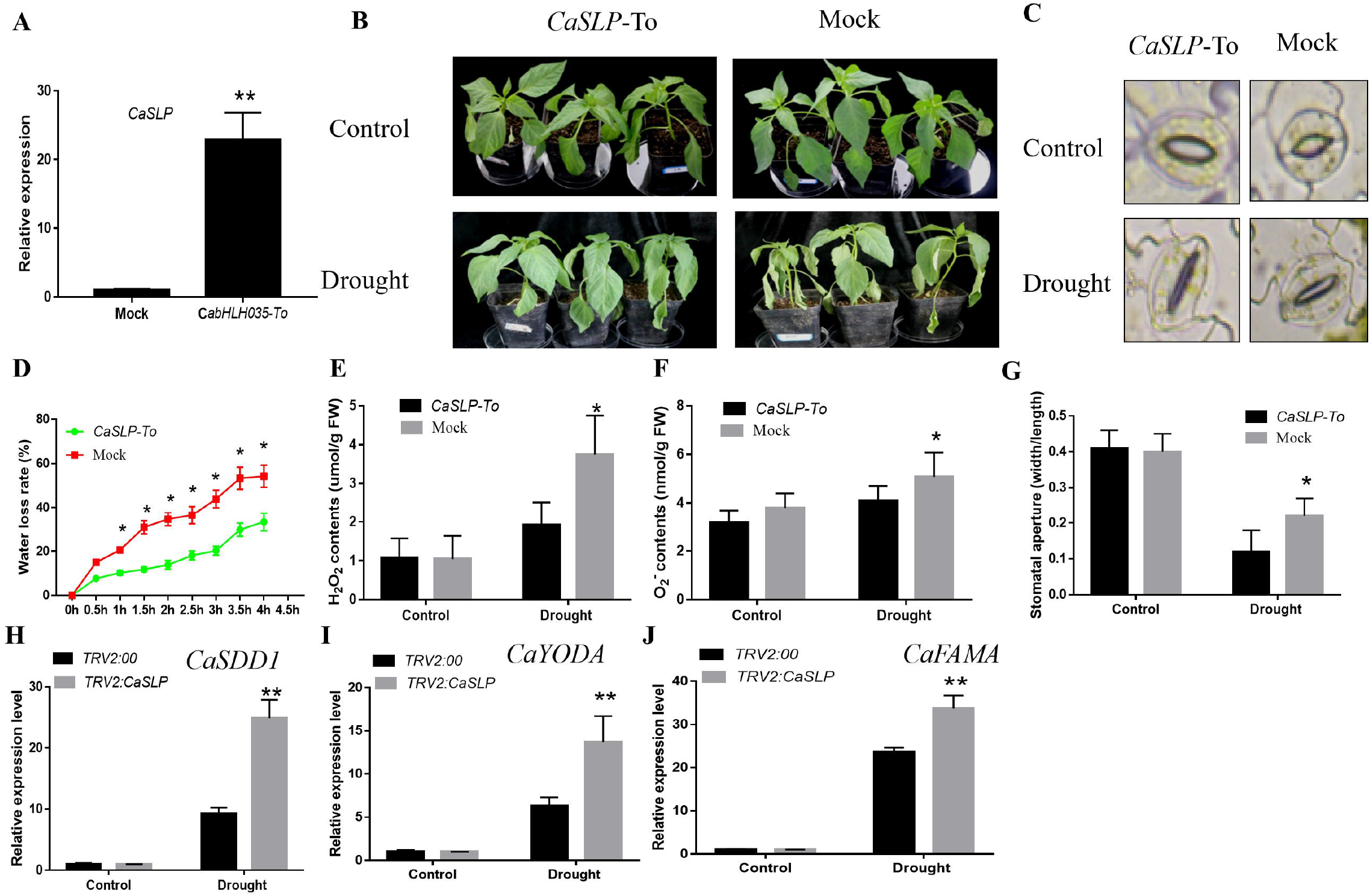
Transient expression of *CaSLP* improved pepper drought tolerance. A, The transcript levels of *CaSLP*. B, Phenotypes of *CaSLP*-To and the control plants under drought stress. C, G. Stomatal aperture of *CaSLP*-To and the control plants under drought stress. D, Water loss rate. E, F. H_2_O_2_ and O2.- contents of *CaSLP*-To and the control plants under drought stress. H-J. The transcript levels of stomatal development-related genes including *SDD1, YODA* and *FAMA* in pepper. The data represent the means ± SDs (n=3, *P < 0.05, **P <0.01, Student’s t-test).

### 3.4 Overexpression of *CaSLP* in Arabidopsis increased drought tolerance

To analyze the functions of *CaSLP* in drought tolerance, three transgenic lines of *CaSLP* were selected, named #1, #2 and #3. Under normal conditions, there were no apparent difference between the *CaSLP* transgenic and the wild type (WT). After drought stress, the WT showed more severe leaf wilting than the *CaSLP* transgenic lines (Figure 4A, B). Better growth performance of the *CaSLP* transgenic lines was supported by Relative Electrolyte leakage (REL), malondialdehyde (MDA) and chlorophyll contents, after upon the drought exposure, the *CaSLP* transgenic lines displayed lower REL and MDA content, and higher chlorophyll content than WT (Figure 4C-E). These results suggest that overexpression of *CaSLP* in Arabidopsis promoted drought tolerance.

**Figure 4.**
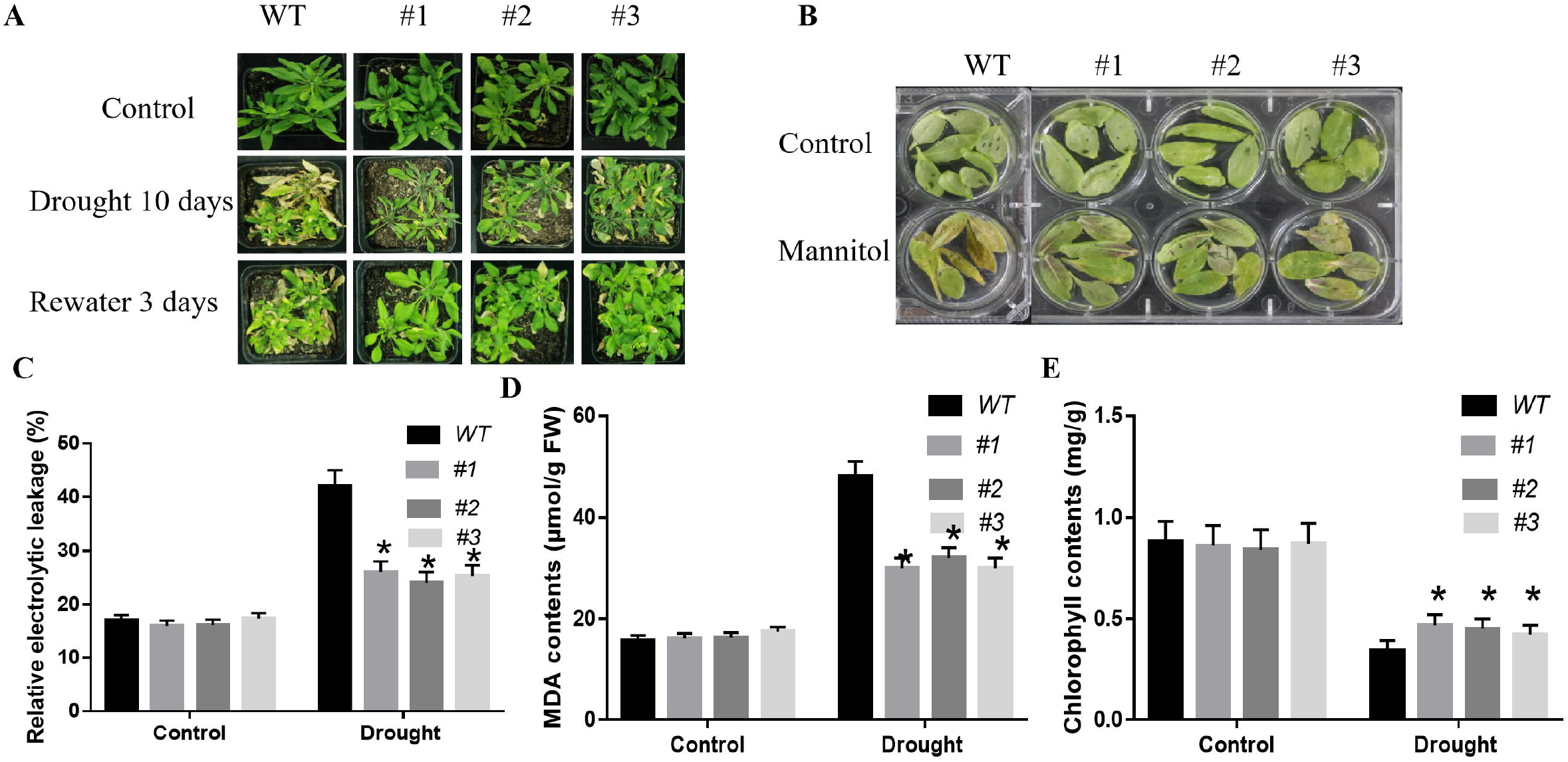
*CaSLP* overexpression in Arabidopsis enhances resistance to drought stress. A, B. Phenotypes of transgenic Arabidopsis and WT under drought stress. C, REL. D, MDA contents. E, Chlorophyll contents. The data are means ± SDs (n=3, *P < 0.05, **P <0.01, Student’s t-test).

### 3.5 Exogenous spring salicylic acid enhanced drought tolerance of *CaSLP*-silenced pepper plants

The obvious induction of *CaSLP* in response to drought (Fig. 1), we conjecture *CaSLP*-mediated drought tolerance via salicylic acid pathway. Therefore, the *CaSLP*-silenced pepper and control plants were sprayed exogenously 2 mM salicylic acid (Fig. 5A). Without stressful conditions, there were no obvious changes of proline, REL, and MDA contents between the *CaSLP*-silenced and control plants. Whereas after exogenous spring salicylic acid, the *CaSLP*-silenced pepper plants exhibited lower REL, MDA, and lower proline contents in comparison with the plants that treated with water interestingly (Fig. 5B-D). These above mentioned data demonstrate that exogenous application of salicylic acid drastically enhanced drought tolerance, further showed the importance of salicylic acid in *CaSLP*-regulated drought tolerance.

**Figure 5.**
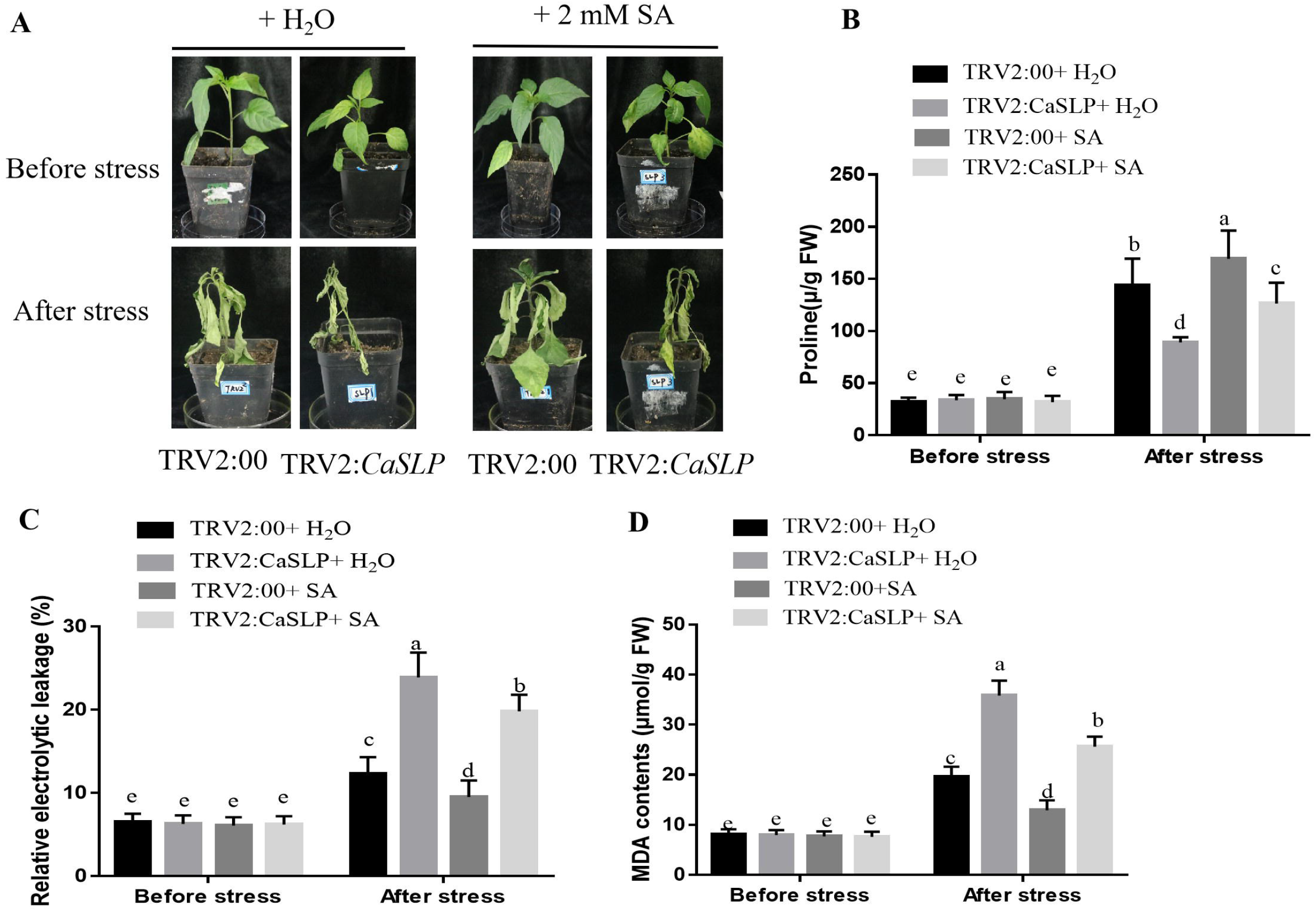
Exogenous spring Salicylic Acid enhanced drought tolerance. A, Phenotypes of *CaSLP*-silenced and the control plants. B, Proline contents. C, REL. D, MDA contents. The data are means ± SDs (n=3, *P < 0.05, **P <0.01, Student’s t-test).

### 3.6 Altered expression of stomatal development-related genes

Next, we measured stomatal development-related genes in pepper, including *SDD1, YODA* and *FAMA*. The qRT-PCR results showed that expression of *SDD1, YODA* and *FAMA* was decreased in *CaSLP*-silenced than control (Fig. 6A-C), suggesting that *CaSLP* acts as a positive regulator of drought signaling. We also measured the expression level of stomatal development-related genes in Arabidopsis (*AtSDD1, AtYODA, AtFAMA, AtTMM* and *AtMPK3*) after exposure to drought stress. The expressions of *AtSDD1, AtYODA, AtFAMA, AtTMM, AtMPK3* and *AtMPK3* were obviously greater in the transgenic plants as compared to the WT plants (Fig. 6). These results indicate that overexpression of *CaSLP* increases drought tolerance in Arabidopsis.

**Figure 6.**
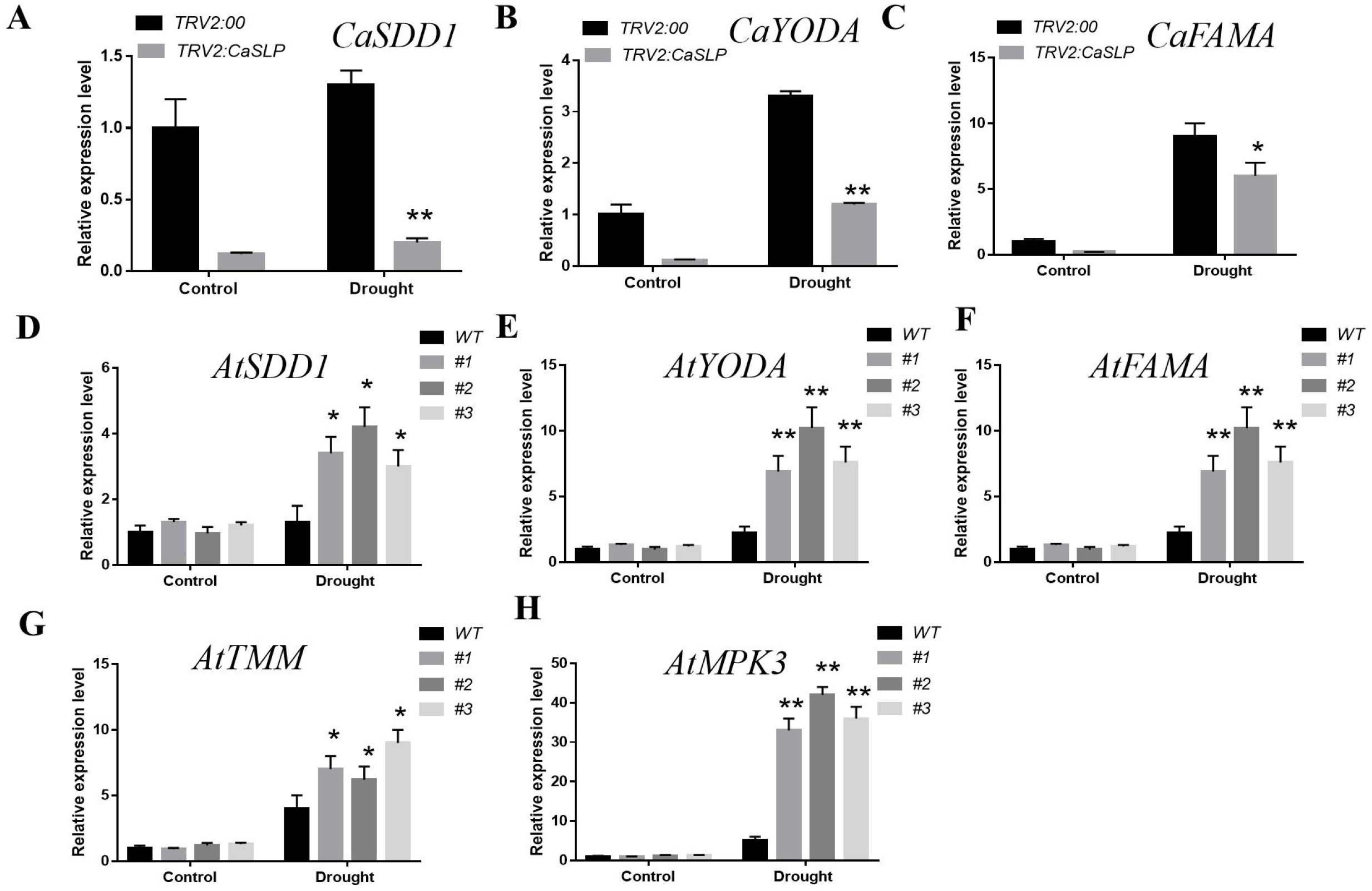
Assessment of the transcript levels. A-C. The transcript levels of stomatal development-related genes including *SDD1, YODA* and *FAMA* in pepper, D-H. *AtSDD1, AtYODA, AtFAMA, AtTMM, AtMPK3, AtPAL3, AtICS* and *AtNPR1 in* Arabidopsis. The data are means ±SDs (n=3, *P < 0.05, **P <0.01, Student’s t-test).

### 3.7 *CaSLP*-knockdown pepper plants demonstrated decreased *Pst.DC3000* tolerance

To examine the *Pst.DC3000* sensitivity, the *CaSLP*-silenced plants and control leaves were infected with *Pst.DC3000*. After *Pst.DC3000* treatment, the leaf color of *CaSLP*-knockdown plants showed more yellow and wilted than control. As indicated by the lower chlorophyll contents of *CaSLP*-knock down plants (Fig. 7A, E, H). Additionally, to explore the accumulation of reactive oxygen species (ROS) post *Pst.DC3000* stress in the control and silenced pepper plants, the Trypan blue and DAB staining of the leaf samples were performed. Figure 7B, F showed that as compared to control, the accumulation of H_2_O_2_ and O_2_^.-^ contents of *CaSLP*-silenced plants was markedly higher than the control plants (Fig. 9I, M). These results showed that the *CaSLP*-silenced plants had more ROS accumulation than WT and *CaSLP*-knockdown pepper plants demonstrated decreased *Pst.DC3000* tolerance, as revealed by reduced cell death and bacterial numbers in pepper leaves compared to the control plants (Fig. 7C, G). To investigate the role of *CaSLP*-knock down pepper plants, the bacterial numbers of *CaSLP*-silenced and control plants was measured. The bacterial numbers of *CaSLP*-silenced showed higher control plants after *Pst.DC3000* treatment (Fig. 7D). Next, we measured the expression of stress related genes *CaNPR1, CaABR1* and *CaPR1*. Further analysis showed the relative expression of *CaNPR1, CaABR1* and *CaPR1* in *CaSLP* VIGS plants were strikingly lower than the controls pepper plants (Fig. 7J-L). Collectively, these results illustrated that *CaSLP* plays a significant role in response to *Pst.DC3000*.

**Figure 7.**
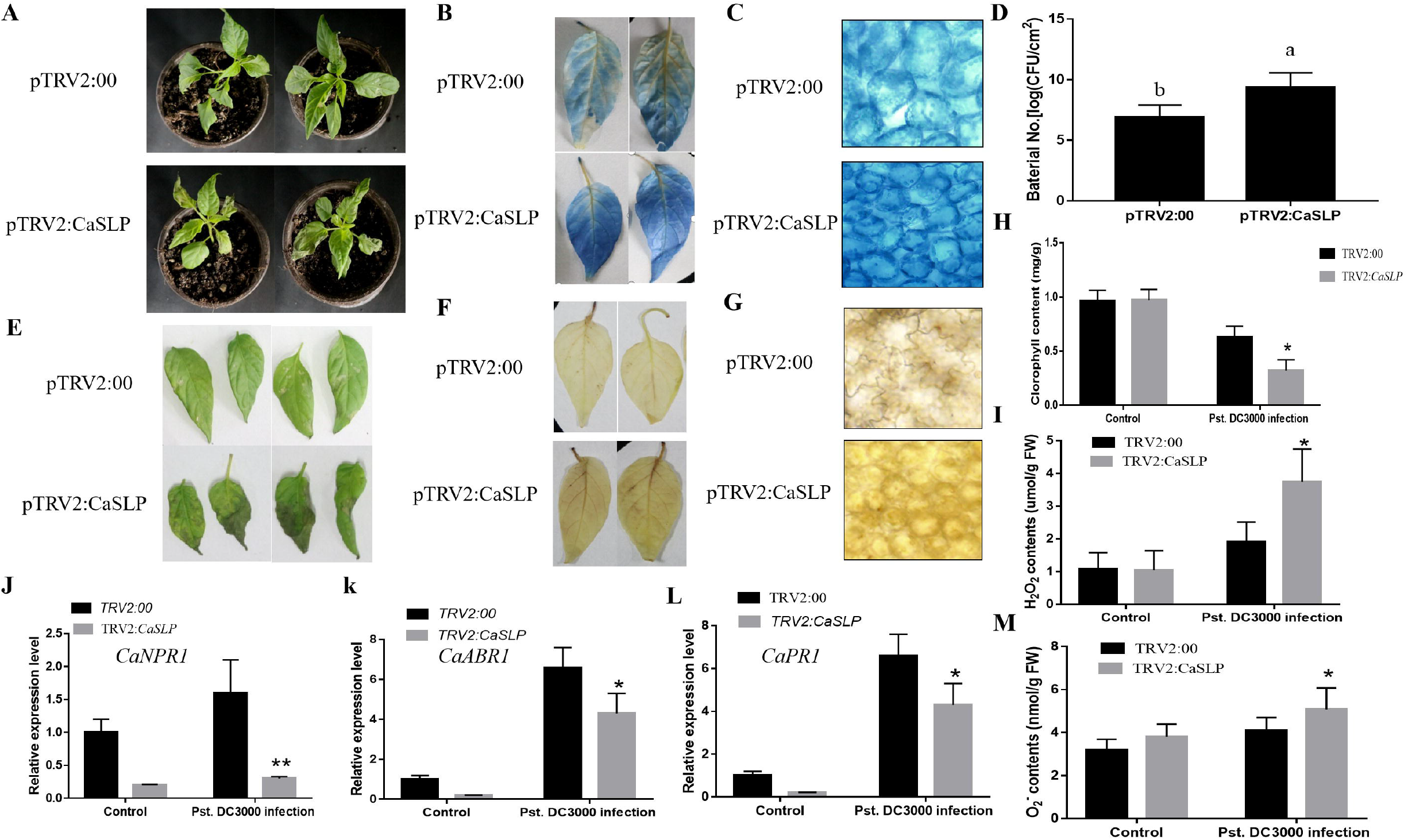
Assessment of *CaSLP*-silenced plants in response to *Pst.DC3000* stress. A, E The phenotypes of *CaSLP*-silenced and the control plants. B, F. Trypan blue staining and DAB staining. C, G Trypan blue and DAB staining for cell death. D, Bacterial numbers. H, Chlorophyll contents. I, H_2_O_2_ contents. J-L, The expression of SA response genes of *CaNPR1, CaABR1* and *CaPR1*. M, O_2_^.-^ content. The data are means ± SDs (n=3, *P < 0.05, **P <0.01, Student’s t-test).

**Figure 8.**
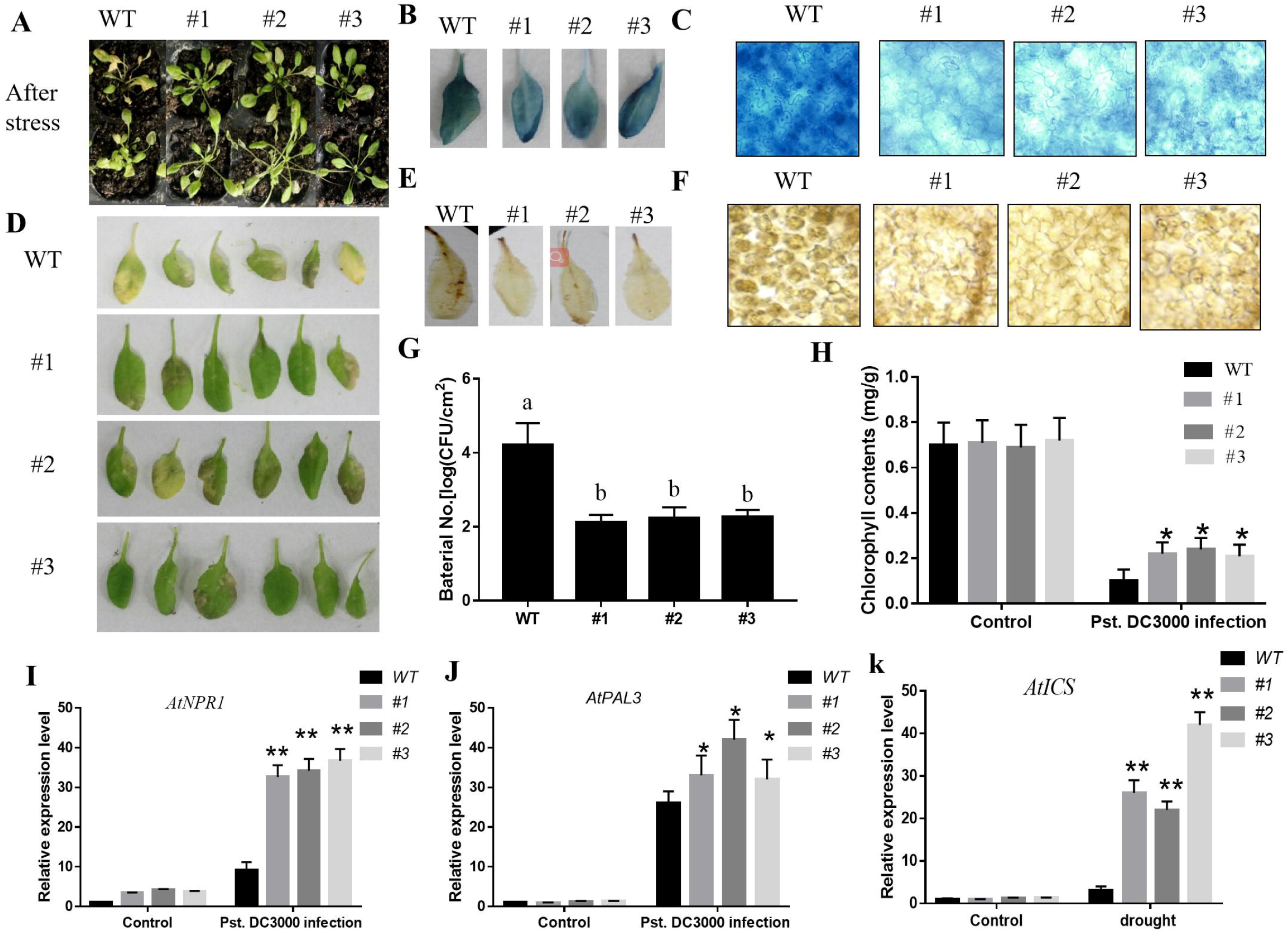
*CaSLP* overexpression in Arabidopsis enhances resistance to *Pst.DC3000* stress. A,D Phenotypes of transgenic Arabidopsis and WT under *Pst.DC3000* stress. B, E Trypan blue staining and DAB staining. C, F, Trypan blue and DAB staining for cell death. G, Bacterial numbers. H, Chlorophyll contents. I-K, The expression of SA response genes of *AtNPR1, AtPAL3*, and *AtICS*. The data are means ± SDs (n=3, *P < 0.05, **P <0.01, Student’s t-test).

**Figure 9.**
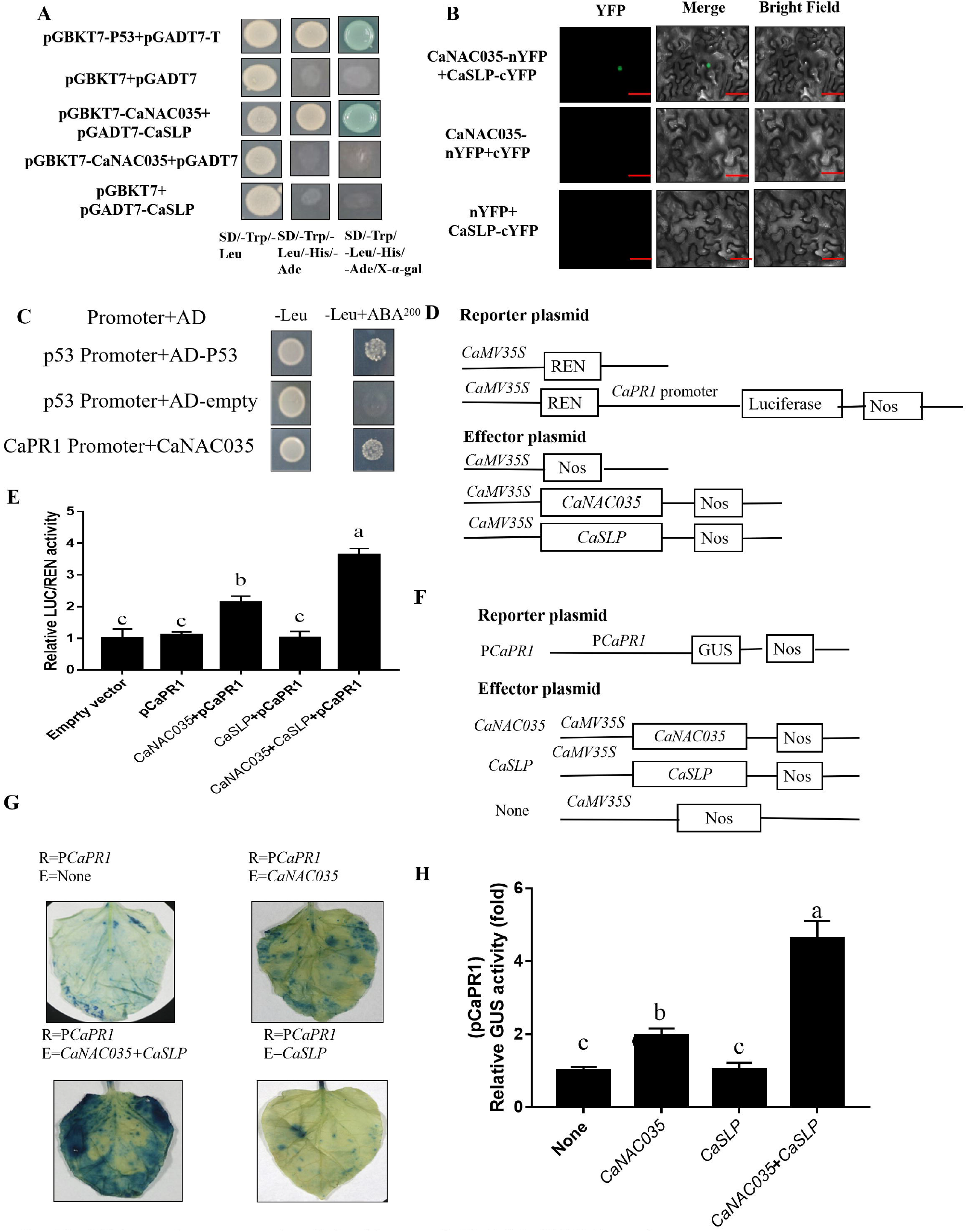
CaSLP enhances the binding of CaNAC035 to its target gene promoters. A, Yeast two-hybrid assay of CaNAC035 and CaSLP. B, Bimolecular fluorescence complementation (BiFC) assay of CaNAC035 and CaSLP. C,Growth of yeast cells. D, F, Diagrams of effector and reporter constructs. E, LUC/REN ratio. G, H, GUS activities. The data are means ± SDs (n=3, *P < 0.05, **P <0.01, Student’s t-test).

### 3.8 Enhanced resistance to *Pst.DC3000* in *CaSLP* transgenic Arabidopsis plants

To explore the role of *CaSLP* in response to disease, the *CaSLP* transgenic lines and WT were infected with *Pst.DC3000*. Under untreated conditions, the color of *CaSLP* transgenic lines and WT was almost the same, there was no substantial difference. Whereas, after 3 days post *Pst.DC3000* injection, the WT showed significant etiolation and yellowing than *CaSLP* transgenic lines, accompanied by WT had lower chlorophyll content than WT (Fig. 8A, D, H). Then, we measured the bacterial numbers of *CaSLP* transgenic lines and *Pst.DC3000*. In the *Pst.DC3000* treatment, the *CaSLP* transgenic lines showed obviously lower bacterial number than WT, respectively (Fig. 8G). Furthermore, we determined the DAB and Trypan blue staining. Under *Pst.DC3000* treatment, the WT showed obviously greater than *CaSLP* transgenic lines, respectively (Fig. 8B, E). these results showed that the transgenic lines had less ROS accumulation than WT and *CaSLP* transgenic lines showed obviously increased resistance to *Pst. DC3000*, as revealed by reduced cell death and bacterial numbers in Arabidopsis leaves compared to WT (Fig. 8C, F). To further understand the underlying molecular mechanism of *CaSLP* in response to disease, we measured the expression level of SA responses and SA biosynthesis related genes *AtNPR1, AtPAL3*, and *AtICS* after *Pst.DC3000* treatment. The expressions of *AtNPR1, AtPAL3*, and *AtICS* were obviously greater in the transgenic plants as compared to the WT plants (Fig. 8I-K). Our results indicated that *CaSLP* transgenic Arabidopsis improved the *Pst.DC3000* tolerance.

### 3.9 *CaSLP* enhances the binding of *CaNAC035* to its target gene promoters

To screen the target protein of CaNAC035, the yeast two-hybrid (Y2H) were performed. To examined the interaction of CaNAC035 with CaSLP, the Y2H and bimolecular fluorescence complementation (BiFC) were used in this study, Y2H results showed that yeast, which containsCaNAC035 and CaSLP grew well on the selective solid medium of SD/-Trp/-Leu/-His/-Ade, and the yeast strains turned blue on the SD/-Trp/-Leu/-His/-Ade/X-a-gal solid medium (Fig. 9A). To further identify the interaction, we performed BiFC assays which showed that co-expression of CaNAC0-nYFP with CaSLP-cYFP displayed significantly fluorescence signaling in the cell nucleus (Fig. 9B). These results showed that CaNAC035 physically interacts with CaSLP in the cell nucleus. The transcripts of *CaPR1* were dramatically increased in the *CaNAC035-*To pepper (Figure S2). We tested whether *CaNAC035* directly regulates *CaPR1* transcription. Typically, NAC TFs could bind to CACG, we analyzed the promoter of *CaPR1* and found that *CaPR1* had a NAC core-binding site. To know the association between CaNAC035 and the *CaPR1* promoter, we carried out Y1H, the results showed that yeast cells transformed including full-length grew well on selective media. (Fig. 9C). These results showed a direct correlation of *CaNAC035* with the promoter of *CaPR1*. To further authenticate the direct binding of *CaNAC035* to *CaPR1* promoter and regulation of expression LUC/REN ratios were performed (Fig. 9D). Dual LUC assays results showed that the *CaPR1* could activate the CaNAC035 expression (Fig. 9E), indicating that *CaNAC035 is* a direct up-stream factor of *CaPR1*. GUS analysis was completed to explore the activity of *CaNAC035pro* with the addition of *CaPR1* (Figure 9F, G, H). These results *CaSLP* enhances the binding of *CaNAC035* to its target gene promoters.

### 3.10 *CaSLP* regulates drought tolerance in a *CaNAC035*-dependent manner

A previous study found that CaSLP interacts with CaNAC035. In order to further verify the relationship between these two proteins, and analyze if *CaSLP* modulates drought stress in a *CaNAC035*-dependent manner, the *CaSLP*-silenced was co-agroinoculated into *35S:CaNAC035:GFP* (Fig. 10A). RT-qPCR showed that expression of *CaNAC035* and *CaSLP* in TRV2:00/35S:*CaSLP*:GFP was significantly than TRV2:*CaNAC035*/35S:*CaSLP*:GFP and TRV2:00/35S:GFP at 12 and 24 h (Fig. 10B, C). Under drought stress, the TRV2:00/35S:*CaSLP*:GFP plants exhibited higher fresh weight and survival rate than TRV2:*CaNAC035*/35S:*CaSLP*:GFP and TRV2:00/35S:GFP plants (Fig. 10D, E). Therefore, the CaNAC035 gene is a necessary for *CaSLP*-mediated of drought stress tolerance.

**Figure 10.**
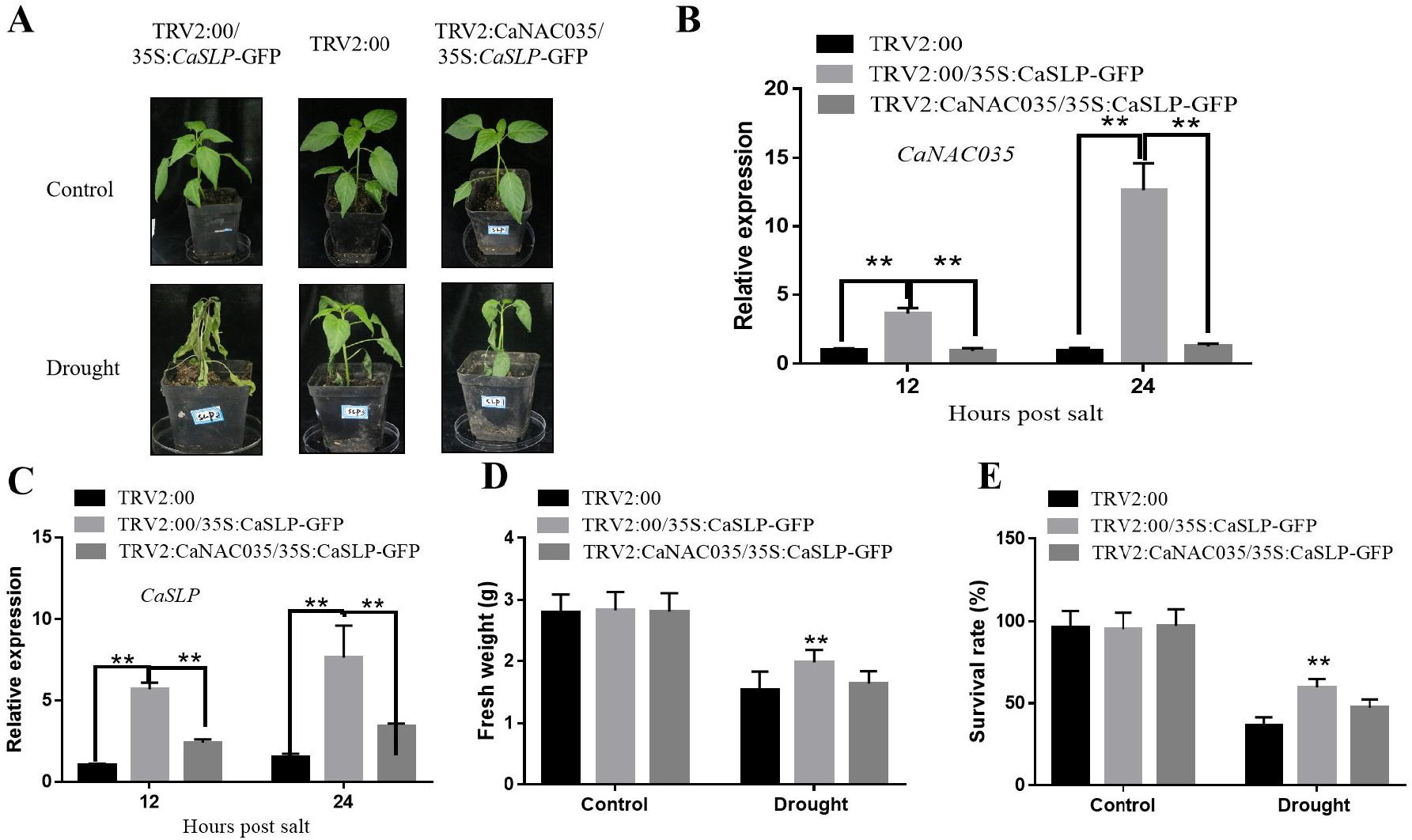
*CaNAC035* is required for *CaSLP*-mediated drought stress tolerance. A, Phenotypes of *CaSLP*-silenced was coagroinoculated into *35S:CaNAC035:GFP*. B, C. The transcript levels of *CaNAC035* and *CaSLP*. D. fresh weight. E. Survival rate. The data are means ± SDs (n=3, *P < 0.05, **P <0.01, Student’s t-test).

## Discussion

Water shortage is a major challenge to agricultural production in the world. Drought stress seriously affects plant growth, development and distribution, resulting in yield decline and economic losses (Kumar et al., 2019). Drought is a critical hazard that influencing the crop yields severity (Sun et al., 2019), nowadays, it’s an important evaluation index to enhance crop drought tolerance (Dunn et al., 2019). In our previous study, we found that *CaNAC035* plays a positive role in stress (cold, salt and drought) tolerance (Zhang et al., 2020). In this study, the Y2H screening was performed, we found that transcription factor CaNAC035 interacted with CaSLP. Further studies have identified that CaNAC035 interacted with CaSLP in the nucleus (Fig. 9B). Sub-cellular location showed that CaSLP localized in the nucleus and cytoplasm (Fig. 1A), the results was consistent with its interaction location in vivo. MDA is considered as a suitable index for lipid per-oxidation when plants are subjected to stress (Mittler et al., 2002). Electrolyte leakage is regarded as an important physiological index to identify stress (Bajji et al., 2002). In this study, the content of REL and MDA in *CaSLP* transgenic lines were significantly lower than WT under drought stress (Fig. 4). In conclusion, these data show that *CaSLP* transgenic lines of lipid per-oxidation and membrane damage may be less in drought condition than in control group.

It is well known that high concentration of reactive oxygen species is toxic to plant cells, which can destroy nucleic acid, oxidize protein and cause lipid peroxidation, and is the main factor affecting cell viability under abiotic stress (Gill and Tuteja, 2010). Drought stress will lead to the accumulation of ROS that adversely affect the growth and development of plants (Miller et al., 2010). ROS had a low level under normal conditions, but when plants are challenged by abiotic stress, ROS can rise sharply, leading to plants death. The accumulation of ROS is used for evaluating the capacity of plants subjected to stress. In this study, the data showed that *CaSLP-*To have less H_2_O_2_ and O_2_^.-^ contents than control after drought stress (Fig. 2). However, *CaSLP*-silenced pepper plants displayed the opposite effects, as shown by histochemical staining (DAB and NBT) (Fig. 3). *CaSLP* contributes to drought tolerance may consistent with the ROS level by regulating the antioxidant gene and maintaining the homeostasis of reactive oxygen species. These results indicated that the transgenic plants exhibited stronger antioxidant capacity than WT, which was related to the significantly decreased ROS level and enhanced antioxidant stress ability of the transgenic overexpression lines, while the VIGS plants showed the opposite trend. Thus, more effective mobilization of the antioxidant system, and thus enhanced antioxidant capacity, is another mechanism by which *CaSLP* exerts a positive effect on drought tolerance.

Stomatal plays a key role in regulating gas and water exchange under stomatal development stage (McKown and Bergmann, 2020). When plants were subjected to drought, the stomatal will be closed preventing water loss (Schroeder et al., 2001). Previously studies have shown that increasing of drought tolerance was connected with the stomatal closure of plants (Aubertet al., 2010). In this study, under drought stress, the *CaSLP*-silenced pepper plants had smaller stomatal apertures compare with control plants (Fig. 2). Our data revealed that *CaSLP* contributes to drought tolerance may involve in stomatal regulation. However, it remains unclear of the regulatory mechanism. These results were consistent with previous reports. For instance, Arabidopsis *AtATAF1* enhances drought tolerance by reducing stomatal aperture (Wu et al., 2009), *AGL16* negatively modulates drought tolerance via stomatal movement in Arabidopsis (Zhao et al., 2020), *AtUNE12* confers salt tolerance by decreasing stomatal aperture in Arabidopsis (He et al., 2022).

SA plays an important role in the plant abiotic stress tolerance, exogenous spring SA can improve plants drought tolerance (Antoni et al., 2016). SA can induce some genes, which were involved in abiotic stresses directly or indirectly (Horváth et al., 2007). SA application slightly enhanced drought tolerance of *CaSLP*-silenced pepper (Fig. 5). Drought stress significantly reduced water loss rate and chlorophyll content, while exogenous salicylic acid had positive effects on these parameters, alleviating the adverse effects of water deficit on plants. Salicylic acid plays an important role in enhancing drought stress tolerance. To further know the molecular mechanisms of *CaSLP* in response to drought stress was attributable to stomatal marker genes, we texted the expression of stomatal marker genes. We found that the stomatal marker genes (*AtSDD1, AtYODA, AtFAMA, AtTMM* and *AtMPK3*) of *CaSLP* transgenic Arabidopsis lines were significantly higher than WT plants (Fig. 6). These data illustrate that *CaSLP* may regulate drought stress responses partially through altering the expression of stomatal marker genes. In this study, we found that the levels of SA response maker genes, including *AtNPR1, AtPAL3*, and *AtICS*. Among the SA response maker genes tested, the *AtNPR1, AtPAL3*, and *AtICS* showed the dramatically increase of expressions in the *CaSLP-OX* than WT (Fig. 8). Heterologous overexpression of CaSLP in Arabidopsis sightly enhanced resistance to the bacterium *Pst.DC3000* via the salicylic acid signaling pathways. However, silencing of *CaSLP* gene resulted in decreased to bacterium *Pst.DC3000* in pepper. Expression levels of SA response maker genes indicated that *CaSLP* may bind to the SA response maker genes promoter, resulting in enhanced *Pst.DC3000* tolerance. There is accumulating evidence indicating that *CaSLP* has important roles in various stresses, it might be related to SA signaling. These data revealed that *CaSLP* roles in response to *Pst.DC3000* stress by participating in the SA signaling pathway.

Based on the Y1H, LUC/REN and GUS results, we found that CaSLP interacts with CaNAC035 and synergistically enhances the transcriptional activity on *CaPR1* (Fig. 9), which indicates its key role in the regulation of stress resistance. The regulatory pathway explains *CaSLP* response to stress tolerance. SA is an important signaling factor in plant stress and involved in some important physiological and biochemical processes in plants. It also plays diverse role in plant stress response in the form of signal molecules, SA plays diverse role in the regulation of resistance to abiotic stress by enhancing the binding ability of SA and its receptor protein, which can perceive and transmit stress signals.

In conclusion, our results found that the transcription factor CaNAC035 interacted with CaSLP in the nucleus, and *CaSLP* plays a positive role in drought stress tolerance in pepper. Herein we proposed the model for *CaSLP* in response to drought and *Pst.DC3000* tolerance stress (Fig. 11).

**Figure 11.**
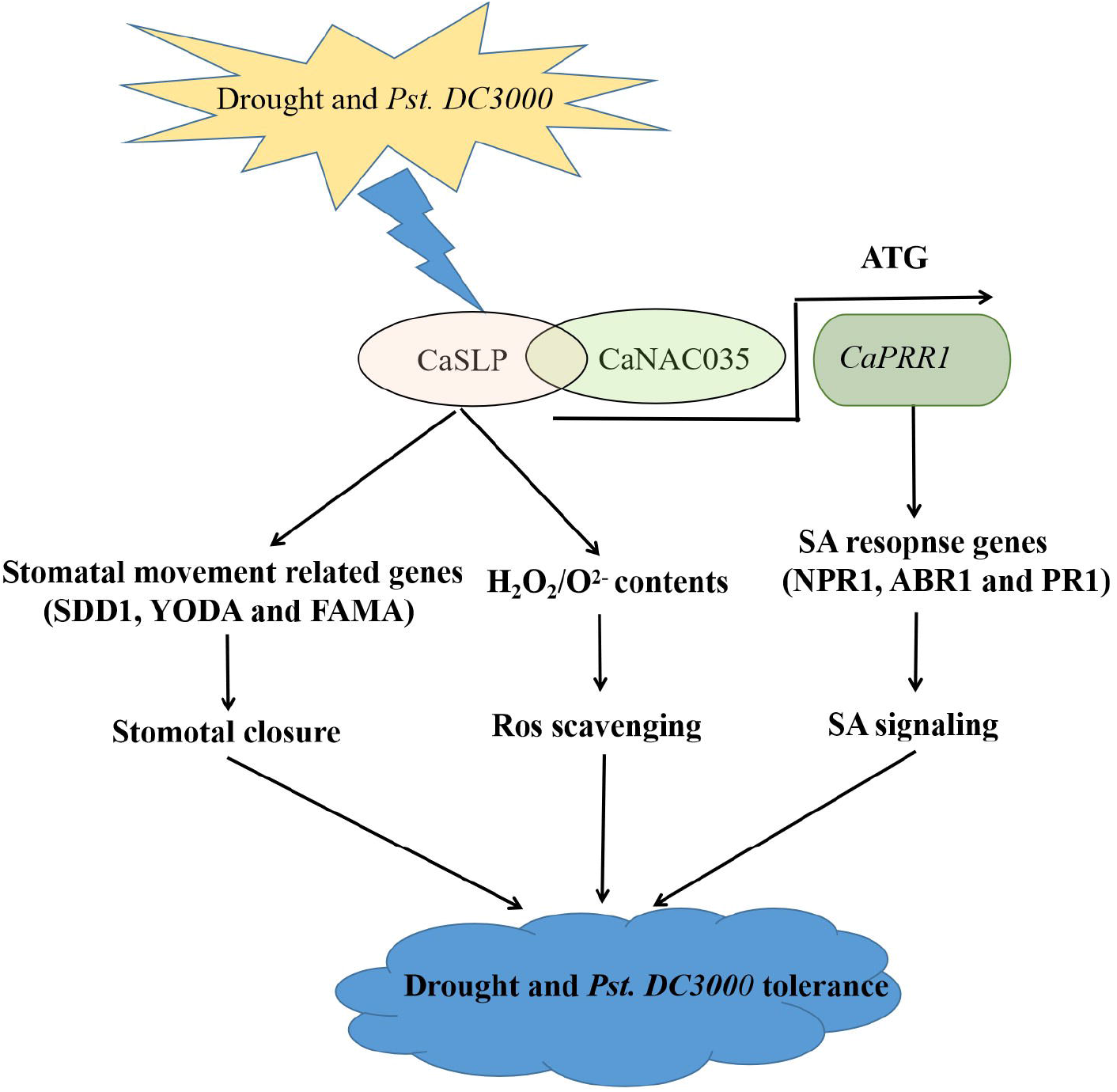
Proposed model for *CaSLP* in response to drought and *Pst.DC3000* tolerance stress.

## Acknowledgements

This research was funded by the National Natural Science Foundation of China (32172582, 316721465), Scientific & Technological Innovative Research Team of Shaanxi Province (2021TD-34), Agricultural Key Science and Technology Program of Shaanxi Province (2021NY-086) and the Natural Science Foundation of Shaanxi Province (2018JM3023).

## Author Contributions

RGC and FHZ conceived and designed the experiments; HFZ, YPP and LC performed the experiments; UQS and KA analyzed the data; FM contributed reagents/materials/analysis tools; HFZ wrote the paper.

## Data availability

The date that support the results that included in this article and its supplementary materials. Other relevant materials are available from the corresponding author upon reasonable request.

## Conflicts of Interest

The authors declare no conflict of interest.

## Notes

### Competing Interest Statement

The authors have declared no competing interest.

